# Copy number signatures in targeted gene panels associate with patient outcomes in routine clinical data

**DOI:** 10.64898/2026.09.20.752979

**Authors:** Emil Netz, Cédric Moris, Amina Hasanovic, Markus Möbs, Thorsten Schlomm, Nikolaj Frost, Raphael Mohr, Sebastian Stintzing, Elena Braicu, Jalid Sehouli, Damian T. Rieke, Manuela Benary, David Horst, Frank Dubois

## Abstract

Targeted gene panels (TGPs) dominate clinical sequencing, yet most copy-number (CN) signature studies rely on genome-wide assays. Whether signatures can be recovered from TGPs and retain clinical relevance remains unclear. We analyzed two real-world TGP cohorts comprising 1,726 patients across 62 tumor types, including 825 with clinical annotations, and made the underlying data publicly available. TGP-derived signatures recapitulated established biological associations, including links to homologous recombination deficiency and TP53 alterations, concordant with published genome-wide assay-derived CN signatures. Using our detailed clinical data, we found TGP-derived CN signatures associated with overall survival under standard therapies in ovarian, pancreatic, and colorectal cancer. In ovarian cancer, CN2 was associated with CCNE1 amplification and shorter survival under paclitaxel/carboplatin, with the survival association validated in an independent SNP-array cohort. These findings demonstrate that routine TGPs yield biologically and clinically relevant CN signatures with potential as biomarkers for therapy stratification.

## 2 Introduction

Recent large-scale whole-genome sequencing-based studies uncovered many potentially clinically impactful alterations in cancer genomes, including signature-based biomarkers predictive of clinical outcomes and therapy response(1, 2). However, despite decreasing sequencing costs, most cancer patients only receive targeted gene panel (TGP) based sequencing of their tumor. In most cancers, this typically results in the analysis of only a handful of specific genetic variants that associate with response to targeted therapies. Whether TGPs retain additional information that could serve to derive signature-based biomarkers remains unclear.

Somatic copy number alterations affect a larger fraction of the cancer genome than any other genetic alteration. Copy number (CN) signatures provide a quantitative framework for inferring the underlying variant-generating processes from sequencing data and have been linked to clinically relevant phenotypes, including homologous recombination deficiency (HRD), chromosomal instability, and therapy response. Given that CN alterations typically span large genomic regions, even the sparse genomic sampling of TGPs may capture sufficient signal. We hypothesized that this property supports CN signature extraction in TGPs.

CN signatures were originally defined in whole-genome sequencing (WGS), whole-exome sequencing (WES), or single nucleotide polymorphism (SNP)-array data (2, 3). These technologies have enabled the discovery of 22 pan-cancer CN signatures, several of which capture well-characterized biological processes—such as the HRD-associated signatures CN17 and CN18. The current COSMIC-anchored CN signature frameworks have primarily focused on linking CN signatures to underlying biological processes, partially due to the lack of robust clinical data for the samples that underwent SNP-array analysis or WGS. Given their much more frequent use in clinical practice, a TGP-based CN signature analysis could allow for their systematic association with clinical variables and outcomes.

The limited understanding of TGP data is also due to the lack of large, fully accessible TGP datasets. While several large TGP cohorts are publicly available, they do not provide allele-specific CN information, raw sequencing, treatment, or survival data (**Table 1**). Consequently, we cannot directly apply existing CN signature methods to these datasets. Two fundamental questions have therefore remained unanswered: (i) whether established CN signatures can be robustly extracted from TGP data despite limited genomic coverage, and (ii) whether TGP-derived CN signatures retain clinically actionable information, such as predicting survival under systemic therapy.

**Table 1:** Publicly available large-scale targeted-panel cancer sequencing cohorts.

| Study/Cohort | Sequenced genes, n | Patients, n | Genome-wide allele-specific CN | Fastq/Bam | Overall survival time | Drug-level treatment | DOI |
| --- | --- | --- | --- | --- | --- | --- | --- |
| Genie | 1 - 760 | 227696 | No | No | Yes | Yes |  |
| MSK-MET | 341 - 468 | 25775 | No | No | No | No | 10.1016/j.cell.2022.01.003 |
| MSK-CHORD | 341 - 505 | 24950 | No | No | No | No | 10.1038/s41586-024-08167-5 |
| Foundation |  |  |  |  |  |  |  |
| Medicine | 287 | 18004 | No | No | No | No | 10.1158/0008-5472.CAN-16-2479 |
| MSK-Impact | 341 - 410 | 10336 | No | No | Yes | No | 10.1038/nm.4333 |
| FoundationOne |  |  |  |  |  |  |  |
| validation | 287 | 2221 | No | Yes | No | No | 10.1038/nbt.2696 |
| Genomic Data |  |  |  |  |  |  |  |
| Commons | 8 - 2.735 | 3373 | Yes | Yes | Partial | Partial |  |
| **Netz et al.** | **56 - 624** | **1726** | **Yes** | **Yes** | **Partial** | **Partial** |  |
| Samstein 2019 | 341 - 468 | 1661 | No | No | No | No | 10.1038/s41588-018-0312-8 |

Here, we present a real-world TGP sequencing dataset of 1726 patients across 62 cancer types, including clinical annotations such as tumor stage and therapy type for 825 patients, which we make publicly available to support broad scientific use. This makes this dataset the only study that provides full access to FASTQs, TGP-based copy number calls, and clinical data aside from GDC (**Table 1**). Using this resource, we demonstrate that CN signatures can be robustly extracted from TGP data and recapitulate known biological associations validated against SNP-array/WGS-based reference datasets. We further show that TGP-derived CN signatures are independently associated with overall survival under standard therapies in three major cancer entities, supporting their potential as clinically accessible biomarkers.

## 3 Results

Our study cohort comprised 1726 cancer patients spanning 62 tumor types, reflecting the diversity of routine molecular diagnostics. The most frequent entities included ovarian (n = 383), prostate (n = 314), colorectal (n = 140), pancreatic (n = 131), and breast cancer (n = 115) (**Fig. 1b**, **Supplementary Table 1**). Of these, 1458 patients were sequenced with the MH Custom Panel V1 (MH; covering 624 genes) and 268 with the North-Eastern German Society for Gynecologic Oncology (NOGGO) panel (56 genes and a CN backbone).

**Figure 1:**
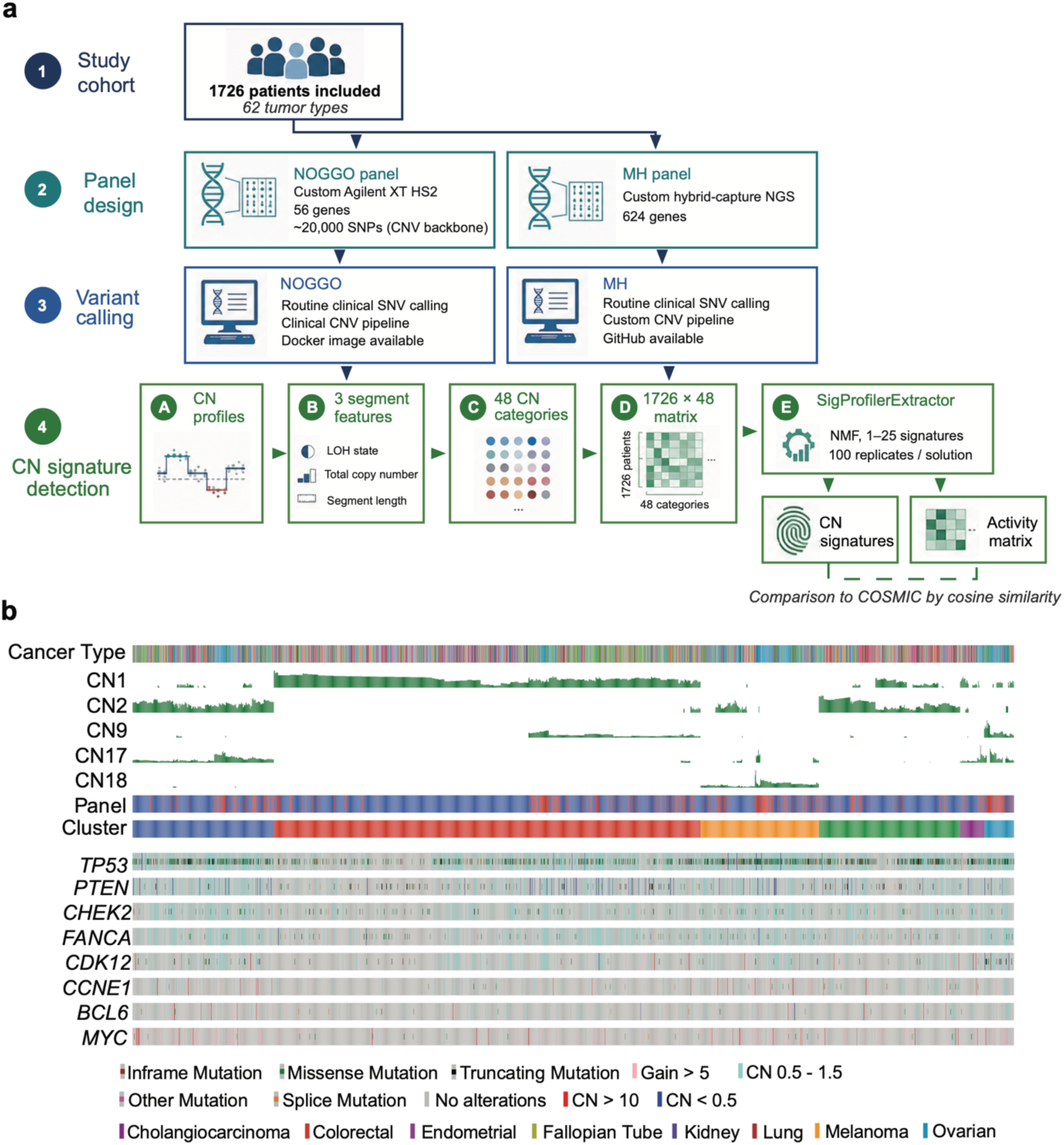
Copy number signatures can be extracted from routinely used targeted gene panels. **a**, CN signature extraction workflow applied to the two targeted gene panels (TGPs) used in this study. **b**, Overview of both TGP cohorts (n = 1C54 samples with activity of at least one of the five CN signatures shared across both panels). Each column is one patient; rows show tumor type, CN signature activity (CN1, CN2, CNS, CN17, CN18), panel (MH/NOGGO), cluster, and selected genetic variants. Patients with only panel-specific CN signatures (e.g., CN3) are shown in Supplementary Fig. 1.

In both panels, we called single nucleotide variants (SNVs) for the covered genes (**Supplementary Table 2**), copy number alterations genome-wide (**Supplementary Table 3)** , and extracted CN signatures using a non-negative matrix factorization framework anchored to the COSMIC reference (**Fig. 1a**)(12). To assess biological validity, we tested for associations between CN signatures and driver-gene alterations and compared our results to a published WGS/SNP-array dataset(2). To evaluate clinical relevance, we analyzed overall survival (OS) in relation to CN signature activity across therapies and tumor types, using clinical data available for 825 patients.

### 3.1 CN signatures are robustly detectable across TGP designs

We identified five CN signatures consistently detected across both panels: CN1 (genomically stable), CN2 (aneuploidy-associated), CN9 (focal LOH), and the HRD-associated signatures CN17 and CN18 (**Fig. 1b**, **Supplementary Table 1**). In the MH panel, we additionally detected CN12, CN13, CN14, and CN16 (**Supplementary Fig. 1a)**. The NOGGO panel cases additionally showed activity of CN3, CN20, and CN11 (**Supplementary Fig. 1b**). These data indicate that established copy number signatures can be detected in commonly used TGPs regardless of the specific panel design.

Unsupervised consensus clustering based on CN signature activity revealed six reproducible clusters comprising samples from both panels, indicating that global CN patterns are not driven by panel-specific biases (**Fig. 1b**, **Supplementary Table 4**). The clusters broadly separated HRD-associated, aneuploid, and genomically stable profiles and showed biologically meaningful tumor-type enrichments. *Cluster 1* combined high CN2 and CN17 activity without significant enrichment for any single entity. *Cluster 2* represented genomically stable samples (high CN1 with some focal LOH/CN9 activity). The cluster was significantly enriched for prostate, endometrial, and cholangiocarcinoma, while showing a significant depletion for ovarian, colorectal, and breast cancer. *Cluster 3* and *Cluster 6* were both associated with ovarian cancer and exhibited high activity of the HRD-related signatures CN17 and CN18, respectively. *Cluster 4* had high CN2 activity and consisted predominantly of colorectal cancer. *Cluster 5* showed low CN17 activity combined with either CN1 or CN2 and was enriched for breast cancer.

The CN signatures extracted from both TGPs showed expected tumor type-specific associations (**Fig. 2a, Supplementary Tables 4-6**). Ovarian cancer was negatively associated with the copy number stable signature CN1 and positively associated with the two HRD-related CN signatures, CN17 and CN18, in both cohorts. The focal LOH-signature CN9 was significantly enriched in prostate cancer. Additionally, CN17 was positively associated with breast cancer in the MH panel, consistent with the known role of HRD in this cancer type(13). These findings demonstrate that TGP-derived CN signatures capture canonical tumor-type-specific genomic processes.

**Figure 2:**
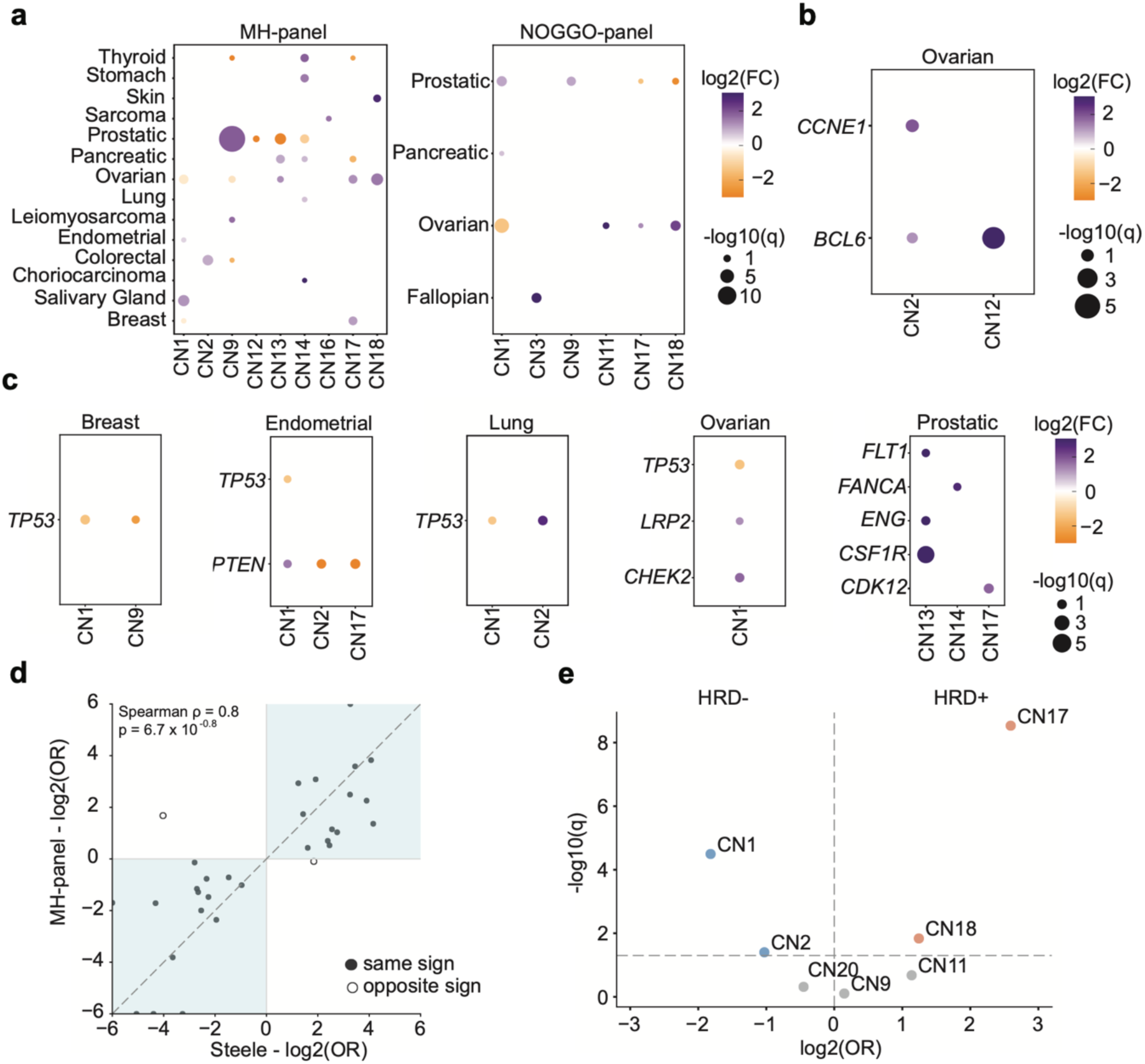
Copy number signatures from targeted gene panels recapitulate tumor-specific genomic features and known biological associations. **a**, Associations between CN signatures and tumor types for MH (left) and NOGGO (right) panels. Dot size reflects significance (-log_10_(q); Mann-Whitney U test), and color indicates effect size (log_2_(FC); purple positive, yellow negative). Only associations with a q value < 0.1 are shown. **b**, Associations between CN signatures and high-level gene amplifications (copy number > 10). Dot size and color are analogous to a. Only significant results are displayed (q < 0.05). Only MH panel results are shown, as no NOGGO associations met the significance threshold. **c**, Associations between CN signatures and driver-gene SNVs in the most common tumor types. Dot size and color are analogous to a and b. Only associations with a q value < 0.1 are shown. Only MH panel results are shown, as no NOGGO associations met the significance threshold. **d**, Concordance of CN signature–SNV associations between the MH panel and a published WGS/SNP-array cohort(2). MH log_2_(OR) (y-axis) vs. reference log_2_(OR) (x-axis); ORs for the MH cohort and the reference were calculated with Fisher’s exact test; filled circles (n = 2S) indicate concordant direction; open circles (n = 2) indicate discordance. Half-circles denote infinite log_2_(OR) values. Only significant (q < 0.05) associations from the reference were included; all MH results are shown regardless of q value. Spearman correlation was used to quantify concordance. **e**, Associations between CN signatures and the GI score (surrogate marker of homologous recombination deficiency, HRD) in the NOGGO cohort. HRD positivity is defined as ≥ 83. Effect size (log_2_(OR), x-axis) and significance (−log_10_(q), y-axis) are shown. CN17 and CN18 show the strongest positive associations, consistent with their HRD-related etiology.

### 3.2 TGP-derived CN signatures recapitulate associations with genetic variants and variant-generating processes previously defined in higher genomic coverage-based studies

We next evaluated whether CN signatures derived from TGP data recapitulate known associations with genetic alterations (**Fig. 2b-e** and **Supplementary Tables 4-6**). This analysis revealed the expected negative association between CN1 and *TP53* mutations across multiple tumor entities (**Fig. 2c**). The aneuploidy signature CN2 was positively associated with *TP53* mutations across pan-cancer and reached significance in the lung cancer cohort. CN17 was positively associated with *CDK12* mutations in prostate cancer, a gene that is known to be involved in the emergence of HRD(14).

Comparison with a published WGS/SNP-array dataset(2) demonstrated strong concordance (29/31 associations in the same direction; Spearman ρ = 0.8, p = 6.7 x 10^-8^; **Fig. 2d**).

CN2 activity was significantly enriched in ovarian cancers harboring high-level amplifications (copy number > 10) of *CCNE1* (n = 6, log_2_(FC) = 1.91, q = 0.02) and *BCL6* (n = 10, log_2_(FC) = 3.88, q = 1.35 x 10^-5^; **Fig. 2b**) in the MH cohort, linking this signature to known drivers of therapy resistance(15, 16).

Genomic instability (GI) scores, available for the NOGGO cohort as a surrogate for HRD(4, 17), also supported these findings. CN17 showed a strong positive association with GI positivity (OR = 6.03, q = 2.93 × 10⁻⁹), whereas CN1 was negatively associated (OR = 0.28, q = 3.2 × 10⁻⁵; Fig. 2e).

Together, these findings suggest that TGP-derived CN signatures closely resemble CN signatures from higher-coverage platforms in their associations with genetic variants and the GI score.

### 3.3 TGP-derived CN signatures independently stratify overall survival under standard systemic therapies in three major cancer entities

We next assessed the clinical relevance of CN signatures using multivariate Cox regression models adjusted for age and tumor stage, within entity-therapy subgroups. Of 825 patients with clinical data, 688 were from the MH cohort and 137 from the NOGGO cohort. In ovarian cancer treated with Paclitaxel and Carboplatin (the current standard of care(18)) CN2 activity was independently associated with shorter overall survival in the MH cohort (n = 57, HR per unit activity = 1.03, 95% CI = 1.01 – 1.05, q = 0.02; **Fig. 3a).** This association was also observed in binary classification when considering any activity of CN2 as “CN2-Active” (n = 57, HR = 2.5, 95% CI = 1.28 – 4.88, q = 0.14; **Supplementary Fig. 2a**). Kaplan-Meier analysis also showed inferior survival in CN2-active versus CN2-inactive patients (n = 57, log-rank p < 0.05; **Fig. 3b**). The association did not reach significance in the NOGGO cohort (**Supplementary Table 6**), likely due to the significantly shorter median follow-up time in this cohort (MH: 41.4 months, NOGGO: 18.3 months). The survival divergence between the CN2-positive and CN2-negative subgroups in the MH cohort emerged only after 2 years. Restricting analysis to patients with > 2.5 years of follow-up in the NOGGO cohort revealed a consistent directional trend (HR = 1.29, 95% CI = 0.94 – 1.77, p = 0.11; **Fig. 3c**).

**Figure 3:**
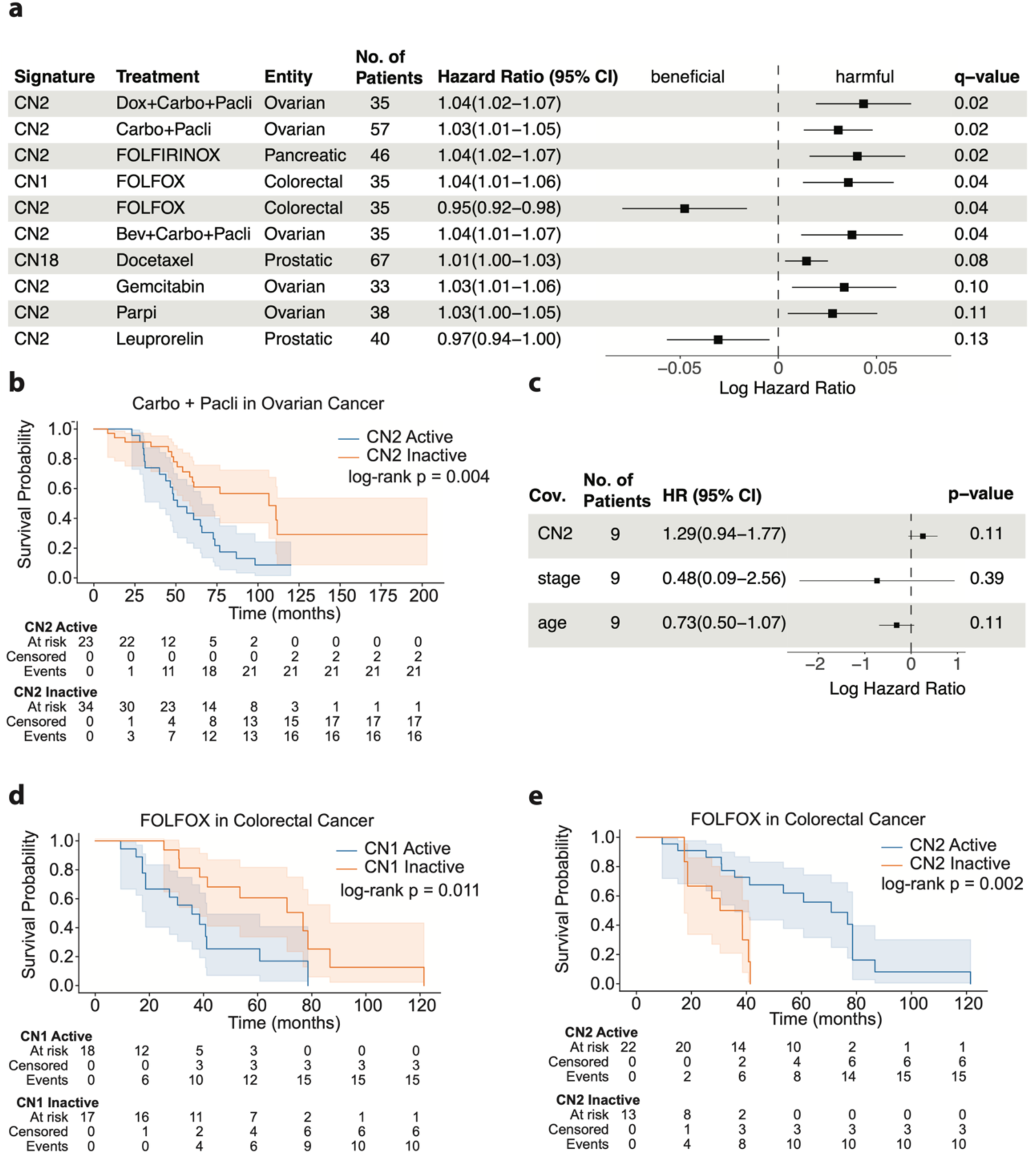
CN signature associations with therapy response. **a**, Multivariate Cox proportional hazards analysis of overall survival (OS) across tumor–therapy subgroups in the MH cohort (n = C88), adjusting for CN signature activity, age, and tumor stage. Therapies with >80% co-administration were merged. Hazard ratios (HRs) per unit increase in CN signature activity are shown with S5% confidence intervals. Only q values < 0.15 are shown; age and stage HRs omitted for clarity. **b**, Kaplan-Meier curves for OS in CN2-active (CN2 > 0) versus CN2-inactive ovarian cancers treated with Carboplatin and Paclitaxel (n = 57). P-value from two-sided log-rank test. **c,** Survival analysis in the NOGGO Carboplatin and Paclitaxel treated ovarian cancer cohort analogous to a, restricting to cases with >2.5 years of follow-up (n = S; median follow-up 18.3 vs. 41.4 months in MH); the CN2–OS association shows a similar trend but does not reach significance. **d**, Kaplan-Meier curves for OS in CN1-active versus CN1-inactive colorectal cancers treated with FOLFOX (n = 35). **e**, Kaplan-Meier curves for OS in CN2-active versus CN2-inactive colorectal cancers treated with FOLFOX, showing an opposing pattern to CN1 (n = 35). Survival curves were compared using the two-sided log-rank test.

We further evaluated this association using SNP-array-derived CN signatures of 485 ovarian cancers in the GDC cohort. Among patients treated with paclitaxel and carboplatin and with available survival data (n = 371), CN2 activity was associated with shorter overall survival in Kaplan–Meier analysis (log-rank p = 0.035). A consistent association was observed in Cox regression adjusted for age and tumor stage, although it did not remain significant after multiple-testing correction (HR = 1.32, 95% CI = 1.01–1.71, q = 0.33; **Supplementary Fig. 3a, c**). These results provide directional external support for the association observed in the MH cohort.

In the GDC cohort, CN20 activity was additionally associated with shorter overall survival among ovarian cancer patients treated with docetaxel and carboplatin, an association that was not detected in our targeted-panel cohort (**Supplementary Fig. 3a, b**). CN20 is characterized partly by relatively short segments with copy-number states of three to four. We therefore hypothesize that the lower genomic resolution of targeted gene panels may cause some of these segments to be merged or assigned to broader aneuploid segments, potentially reducing the distinction between CN20 and CN2 (**Supplementary Fig. 3d**). This interpretation remains to be formally evaluated.

Together, these findings nominate CN2 as a candidate biomarker for Paclitaxel/Carboplatin resistance in ovarian cancer.

In pancreatic cancer treated with the FOLFIRINOX regimen, CN2 activity was also associated with significantly shorter overall survival (HR = 1.04, 95% CI = 1.02 – 1.07, q = 0.02, **Fig. 3a, Supplementary Fig. 2b**), suggesting that the prognostic impact of aneuploidy-associated CN states extends beyond ovarian cancer. In colorectal cancer treated with FOLFOX, CN1 activity was associated with inferior overall survival (HR = 1.04, 95% CI = 1.01 – 1.06, q = 0.04, **Fig. 3d**), while CN2 activity was associated with improved overall survival (HR = 0.95, 95% CI = 0.92 – 0.98, q = 0.04, **Fig. 3e**). For both pancreatic and colorectal cancer, no sufficiently large cohorts treated with these regimens were available in the GDC (**Supplementary Table 7**). These therapy-specific patterns indicate that CN signature activity may stratify outcome differently depending on both tumor entity and treatment context.

## 4 Discussion

Here, we show that CN signatures can be robustly extracted from targeted gene panel data and retain both biological and clinical relevance. Despite limited genomic coverage, TGP-derived signatures recapitulate key features previously described in SNP-array and WGS-based studies(2, 3, 7, 19, 20). This provides evidence that much of the information needed for CN signature extraction is already present in routinely used TGPs. We also found associations between the TGP-based CN signatures and therapy response in ovarian, pancreatic, and colorectal cancers, independent of clinical covariates, suggesting these signatures as potential biomarkers.

A central finding is the consistent association of the aneuploidy-related CN2 signature with adverse outcomes in platinum-treated ovarian cancer. This observation is mechanistically supported by the enrichment of CCNE1 (Cyclin E1) amplification. Cyclin E1 is involved in G1/S cell cycle transition, and its overexpression causes replication stress and genomic instability(21). Multiple studies have associated CCNE1 amplification with primary chemotherapy resistance in ovarian cancer (15, 16). However, the exact copy number cutoff of *CCNE1* above which that effect becomes clinically relevant remains unclear. *CCNE1* amplifications were reported to be significantly depleted in HR-deficient ovarian cancers(22, 23). Mechanistically, *CCNE1* amplification was shown to lead to aneuploid tumors with functional HR pathways(24, 25). This mechanism could explain both the decrease in platinum agent efficacy and the increase in CN2 activity. *PKMYT1* and *ATR* inhibitors(26) have been shown to induce synthetic lethality in *CCNE1*-amplified ovarian and endometrial cancer xenografts. Overall, while CCNE1 amplification was repeatedly associated with resistance in ovarian cancer, the precise genomic context in which it becomes clinically relevant remained incompletely defined. Our data suggest that CN2 may capture a resistant, aneuploid ovarian subgroup that is not currently identified in routine diagnostics.

We also found CN2 linked to poor survival in pancreatic cancer treated with FOLFIRINOX. The effects of aneuploidy on therapy response remain understudied in this cancer type. However, a recent study showed that the group 2 tandem duplication phenotype (TDP) was associated with worse survival in advanced-stage pancreatic ductal carcinoma treated with platinum-based therapy(27). The group 2 TDP is characterized by HR proficiency, CCNE1-amplification, and a larger tandem duplication span size compared to group 1 TDP(27, 28). As CN2 captures large-scale CN alterations, it is plausible that it reflects a similar genomic context. Consistent with our observations in ovarian cancer, CN2 activity may therefore be enriched in HR-proficient, aneuploid pancreatic tumors with reduced sensitivity to platinum therapy.

Copy number signatures reflect entire mutational processes instead of single genetic variants. Utilizing copy number signatures to detect HRD, for example, can thus lead to higher sensitivity compared to conventional scores(29). Therefore, using CN2 as a surrogate for *CCNE1*-driven chemotherapy resistance might capture mechanisms previously undetectable from TGPs, such as enhancer amplifications(30).

Our findings also align with the broader literature on CN signatures and treatment response. Recent studies have shown that chromosomal instability-related signatures can predict benefit from taxane, platinum, and anthracycline therapy in ovarian cancer(31). That study also demonstrated concordance between WGS and TGP CN signatures in eight cases using platform-specific mixture-model approaches. Our study extends these findings to 1,726 patients across 62 tumor types. In contrast to platform-specific mixture-model approaches, the COSMIC-anchored framework used here supports cross-platform comparison and facilitates broader clinical implementation(2).

Several studies have demonstrated links between CN signature activity and clinical outcomes(3, 7, 19, 20). Using the detailed treatment history available for a subset of patients in our study, we additionally found that CN2 identifies a subgroup of ovarian cancers that potentially benefits less from Paclitaxel/Carboplatin treatment. Our data also suggest that the association between CN signatures and clinically relevant outcomes could vary depending on tumor type and treatment context. More data will be needed to develop CN signatures into robust predictive markers, but the possibility of extracting them from widely available TGP data now makes this feasible.

While this study provides evidence for the potential clinical relevance of TGP-based CN signatures, several limitations should be acknowledged. Important prognostic factors, including standardized histological subtypes and residual disease after surgery, were unavailable and could not be included in the Cox models. In addition, many patients received multiple therapies, which required us to combine frequently co-administered treatment regimens in some analyses. This may obscure whether observed effects are attributable to a single drug or a treatment combination. Finally, the shorter follow-up in the NOGGO cohort limited confirmation of some findings observed in the MH cohort.

The MH panel (covering 624 genes) used in this study is comparable in size to widely used assays such as TSO500 (523 genes). In contrast, the smaller NOGGO panel, while covering only 56 genes, incorporates a dedicated copy-number backbone. However, panels lacking either sufficient genomic coverage or a dedicated copy-number backbone may not provide adequate signal for reliable CN signature inference.

Our findings establish CN signature analysis from targeted gene panel data as a feasible and clinically informative approach. Given the widespread use of TGPs, this framework provides a path for CN signature analysis to become a practical extension of existing diagnostic sequencing, with the potential to refine patient stratification and identify therapy-relevant tumor subgroups.

## 5 Methods

### 5.1 Study cohort

1726 patients from the Charité Comprehensive Cancer Center molecular tumor board study (IRB EA1/133/23) were included in this study (**Supplementary Table 1**). All patients provided written informed consent according to the Declaration of Helsinki. The responsible ethics committee at Charité approved the study.

### 5.2 Panel design

#### NOGGO panel

NOGGO is a custom Agilent XT HS2 hybrid-capture next-generation sequencing (NGS) assay (Agilent Technologies, Santa Clara, CA, USA) used for routine molecular diagnostics at Charité. It covers the coding regions of 56 genes and approximately 20,000 selected single-nucleotide polymorphisms (Agilent OneSeq CNV Backbone)(4).

#### MH panel

MH is an Agilent hybrid-capture NGS assay used for routine molecular diagnostics at Charité. It covers the whole exonic region of 624 genes(5).

### 5.3 Variant Calling

#### NOGGO cohort variant calling

For the NOGGO cohort, SNV and CN variant (CNV) information was obtained through a routinely used clinical diagnostics pipeline. SNVs and small indels were called using the SEQNext module of SEQUENCE PILOT (JSI medical systems, Ettenheim, Germany). CNV detection was performed using a custom bioinformatics pipeline based exclusively on publicly available tools, including Mutect and PureCN for allele-specific CN analysis. The Docker image and wrapper code are available on request.

#### MH cohort variant calling

SNV information for the MH cohort was available through routine clinical diagnostics. CNV calling was performed using a custom pipeline (available on GitHub(6)), which integrates publicly available tools such as CNVkit and PureCN.

### 5.4 CN signature detection

#### Extraction

CN signatures were extracted following the framework established by Islam et al., using the SigProfilerExtractor tool (v1.1.24). Briefly, the CN profiles of our 1726 cases (**Supplementary Table 3**) were categorized based on three CN segment-level features (LOH state, total copy number, and segment length), resulting in 48 CN categories. This yielded two non-negative matrices with the dimensions of 1458 x 48 for the MH and 268 x 48 for the NOGGO panel, which were the input for SigProfilerExtractor. To create these matrices from the CN profiles of our samples, we used the read_copynumber function from the sigminer package in R(7).

We ran SigProfilerExtractor with default parameters, performing 100 non-negative matrix factorization (NMF) replicates for each candidate solution with 1 to 25 CN signatures. The tool determines the optimal number of CN signatures by using cluster-based stability metrics across NMF replicates.

Following extraction, SigProfilerExtractor compares CN signatures to the previously reported CN signatures from the COSMIC database by using cosine similarity. In addition to the signatures, NMF yields an activity matrix that quantifies each CN signature’s contribution to each sample.

#### Clustering

In both panels, samples were clustered based on CN signature activity using the ConsensusClusterPlus function implemented in the ConsensusClusterPlus R package. The number of clusters was determined based on the consensus distribution as outlined by Monti et al(8).

#### Tumor type association

The Mann-Whitney U test (mannwhitneyu function from the scipy package in Python) was used to test for associations between the continuous CN signature activity and tumor types. This was done for all CN signature/ tumor type combinations separately in the two panels. All resulting *P* values were corrected using the Benjamini-Hochberg (BH) method.

### 5.5 Signature association with genetic variants and variant-generating processes

#### Driver-gene SNV association

To test for associations between signature activity and driver-gene SNVs, the SNV profiles from 1608 patients (1371 MH, 237 NOGGO) were used (**Supplementary Table 2**). SNVs included missense mutations, small insertions and deletions, and splice-site mutation status. For every gene, we divided our cohort into two groups based on whether an SNV was detected for that gene or not. The Mann-Whitney U test (mannwhitneyu function from the scipy package in Python) was performed between the CN signature activity in the SNV group and the no-SNV group. We also calculated the fold change (FC) by dividing the mean CN signature activity in the SNV group by the mean CN signature activity in the no-SNV group. Additionally, we performed Fisher’s exact test (fisher_exact function from the scipy package in Python) between the described groups for binarized CN signature activity in absent (CN activity = 0) and present (CN activity > 0). This was done separately for both panels and within tumor types for all CN signature/gene combinations. All *P* values were corrected using the Benjamini-Hochberg (BH) method.

To compare the results of this analysis with Steele et al.’s results, we calculated the Spearman correlation between Steele et al.’s statistically significant associations and all MH panel associations (regardless of *q* value).

For a pan-cancer evaluation, we combined *P* values of the gene/CN signature associations using Fisher’s method (combine_pvalues function from the scipy package in Python) if the direction of the association was consistent across all tumor types. The combined *P* values were then corrected using the BH method.

#### Driver-gene CN association

We also tested associations between driver-gene copy number status and CN signatures. Copy number status was defined by four states: Heterozygous loss (CN 0.5 – 1.5), homozygous loss (CN < 0.5), gain (CN 6 - 10), and amplification (CN > 10). As for the SNV associations, the analysis was conducted separately for the two panels and within tumor types, using the Mann-Whitney U test.

#### GI score associations

The GI score quantifies genomic instability and was calculated for all 268 NOGGO cases. It is based on three measures: percent loss of heterozygosity, percent copy number alteration, and percent telomeric copy number alteration(4). We performed a two-sided Fisher’s exact test between signature activity (binarized in present and absent) and the GI score (binarized as ≥ 83 corresponding to HRD positive and < 83 corresponding to HRD negative). *P* values were corrected using the BH method.

### 5.6 Signature association with therapy response

To test for associations between CN signature activity and therapy response, we collected clinical data of 825 cases (688 MH, 137 NOGGO, **Supplementary Table 1**). The cohort was stratified by therapy and entity for the 10 most frequent cancer entities. Within each entity-therapy subgroup, a Cox proportional hazards model was fitted for each CN signature using the CoxPHFitter function from the lifelines package in Python. OS was used as the dependent variable, and CN signature activity (number of CN segments assigned to the signature, continuous: **Fig. 3a**, binary: **Supplementary Fig. 2a**), age, and tumor stage as independent variables.

83% of patients received more than one therapy. To address this, we calculated the fraction of cotreatment for each therapy pair within the entity. If the fraction of co-treatment exceeded 80% for one or more therapies, these therapies were combined for the analysis (e.g., Paclitaxel + Carboplatin in ovarian cancer).

Survival time was defined as the interval from diagnosis to death or censoring. The analysis was performed separately in both TGPs due to the differences in follow-up time. The BH method was used to correct for multiple hypothesis testing. Kaplan-Meier survival curves were generated for results with a *q value* < 0.15 in the multivariate models using the KaplanMeierFitter function from the lifelines package in Python.

### 5.7 Signature analysis in the GDC cohort

To validate findings from the MH and NOGGO cohorts, we analyzed publicly available copy-number and clinical data from 485 patients with ovarian, 390 patients with colorectal, and 141 patients with pancreatic cancer, obtained through the National Cancer Institute Genomic Data Commons (GDC, **Supplementary Table 7**)(9). The analyzed cases originated from The Cancer Genome Atlas (TCGA)(10), the Human Cancer Model Initiative (HCMI), and the Clinical Proteomic Tumor Analysis Consortium (CPTAC)(11). We included only cases with Affymetrix SNP6-array data, allele-specific copy-number calls generated using the ASCAT3 pipeline, and sufficient clinical and treatment information for analysis.

We extracted CN signatures using the procedure described in Section 5.4. We then assessed associations between CN signature activity and therapy response using the approach described in Section 5.6.

## 6. Data availability

Supplementary Table 1 contains clinical information; Supplementary Tables 2 and 3 provide genetic variant information and CN calls. The FASTQ data used in this study are available via the German Human Genome-Phenome Archive (GHGA; data.ghga.de) under the GHGA Accession https://data.ghga.de/study/GHGAS98337713034341. Further details, including the data access policy for the study, can be found there.

## 7. Code availability

All code used for CN signature extraction and association analysis can be found on GitHub at https://github.com/emilnetz/Copy-number-signatures-in-targeted-gene-panels.

## 8. Acknowledgements

FD is supported by the Max-Eder program of the German Cancer Aid (Deutsche Krebshilfe).

## 9. Author contributions

E.N. performed statistical analyses and wrote the manuscript. C.M. generated the allele-specific copy number profiles for the MH cohort. A.H. and M.M. curated the cohorts. T.S., N.F., R.M., S.S., E.B., J.S., D.T.R., and D.H. interpreted and curated data. M.B. supervised C.M. and interpreted data. F.D. initiated and supervised the study and wrote the manuscript. All authors read and approved the manuscript.

## 10. Competing Interests

**R.M.** has received honoraria, research support, and/or travel/accommodation expenses from AstraZeneca, Roche, MSD, Falk, Ipsen, BMS, and Eisai. **S.S.** has received honoraria, research support, and/or travel/accommodation expenses from AMGEN, AstraZeneca, Bayer, BMS, CV6, ISOFOL, Leo-Pharma, Lilly, Merck KGaA, MSD, Pierre-Fabre, Roche, Sanofi, Servier, Taiho, and Takeda. **J.S.** reports grants or contracts from Roche, MSD, GSK, Tesaro, AstraZeneca, Eisai, Merck, and Novocure; consulting fees from Immunogen, Incyte, GSK, AstraZeneca, Clovis, Novocure, MSD, Eisai, and Merck; payment or honoraria for lectures, presentations, speakers bureaus, manuscript writing, or educational events from Immunogen, Incyte, GSK, AstraZeneca, Clovis, Novocure, BMS, Eisai, and Novartis; support for attending meetings or travel from GSK, Astra Zeneca, Roche, Novocure, Immunogen, Incyte, MSD, and Eisai; participation on a data safety monitoring board or advisory board for Immunogen, Incyte, GSK, AstraZeneca, Clovis, Novocure, Bristol Myers Squibb, MSD, Merck, Bayer, and PharmaMar; leadership or fiduciary role in other board, society, committee or advocacy group, paid or unpaid, for ENGAGE, ESGO, ASCO, ESGO, GCIG, Deutsche Stiftung Eierstockkrebs, and AGO; and medical writing assistance from MSD. **D.T.R.** has received honoraria, research support, and/or travel/accommodation expenses from Bayer, Eli Lilly, Bristol-Myers Squibb, Roche, BeiGene, JCJ, and Seagen. The remaining authors declare that they have no competing interests.

---

**Supplementary Figure 1:**
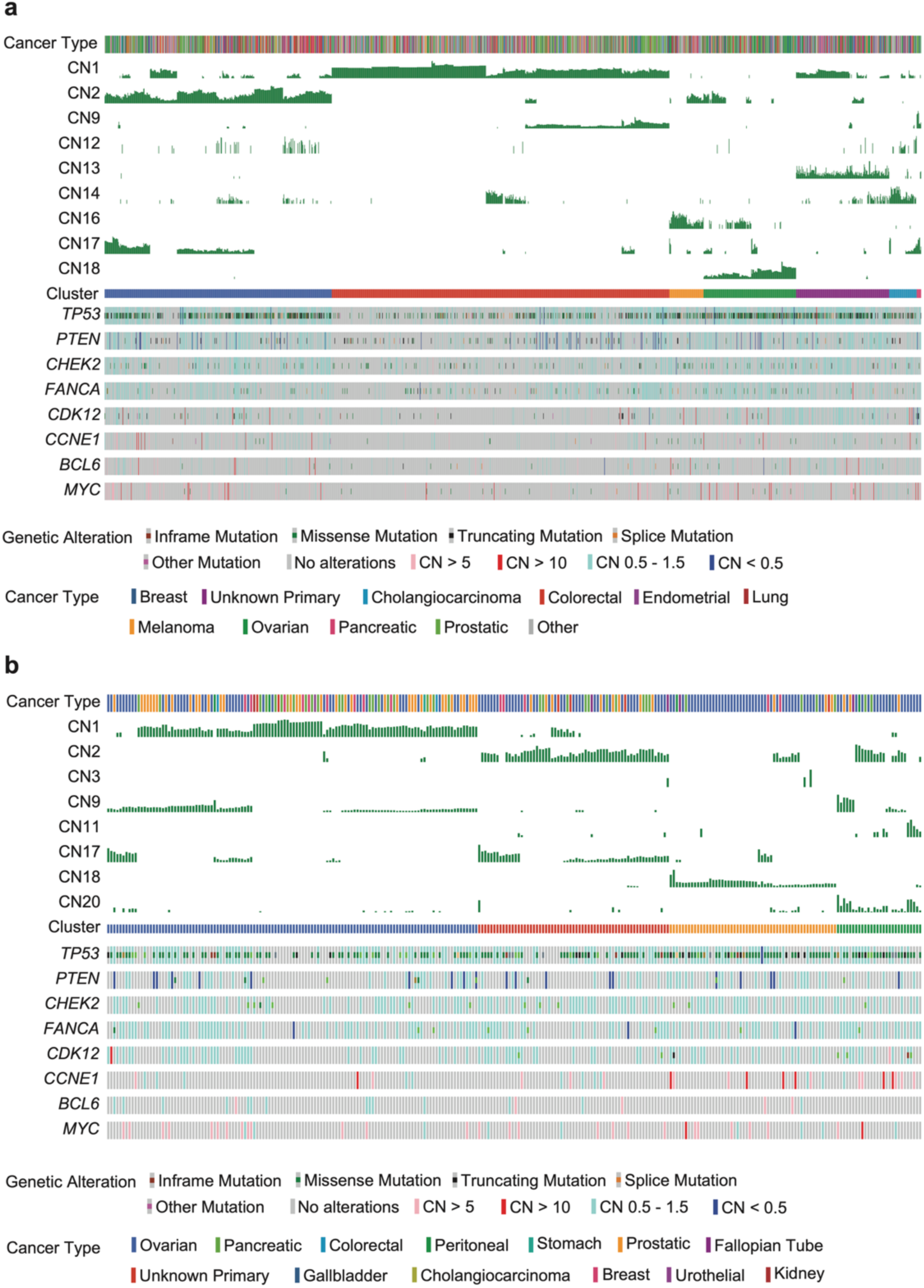
CN signature activity and genetic variants in the MH and the NOGGO cohort. **a**, cBioPortal-style visualization of the MH cohort (n = 1458) showing cancer type, CN signature activity, cluster, and genetic variant per sample. **b**, cBioPortal-style visualization of the NOGGO cohort (n = 268) showing cancer type, CN signature activity, cluster, and genetic variant per sample.

**Supplementary Figure 2:**
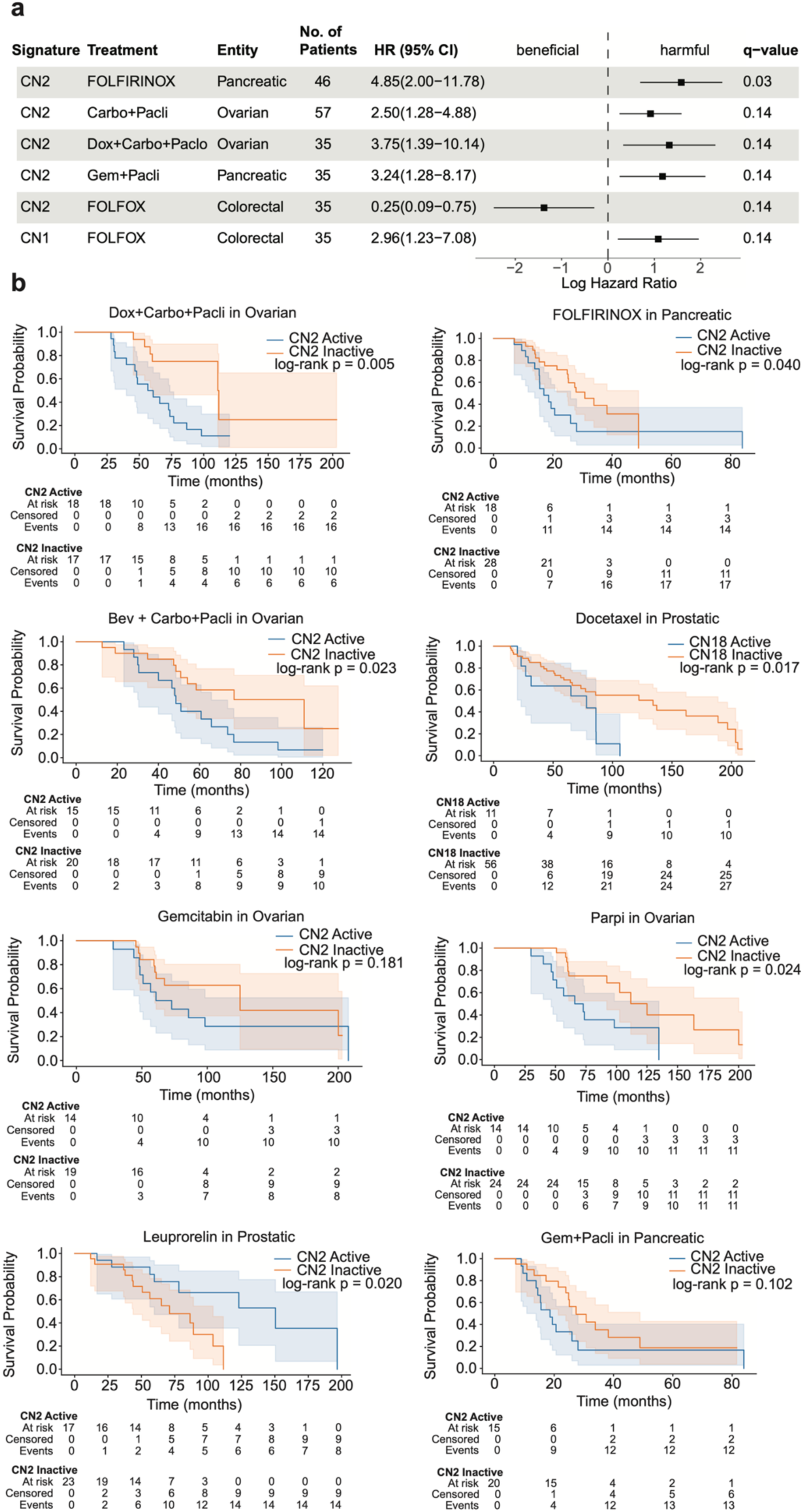
CN signature association with therapy response. **a**, Multivariate proportional hazard models showing the relative risk of CN signature exposures on OS for different therapies and cancer types, controlling for age and tumor stage, in the MH cohort (n = 688). Treatments that regularly co-occur were combined. Contrary to the analysis shown in Fig. 3a, HRs represent the risk of any CN signature activity (CN signature activity > 0) compared to no activity (CN signature activity = 0). Only results with a q value < 0.15 are shown. HR of age and tumor stage are not shown. **b,** Kaplan-Meier curves comparing OS of CN-active (CN signature activity > 0) vs CN-inactive (CN signature activity = 0) cases for different tumor types and therapies. The selection of curves shown is based on the results of the multivariate proportional hazard models (Fig. 3a, Supplementary Fig. 2a). Survival curves were compared using the two-sided log-rank test.

**Supplementary Figure 3:**
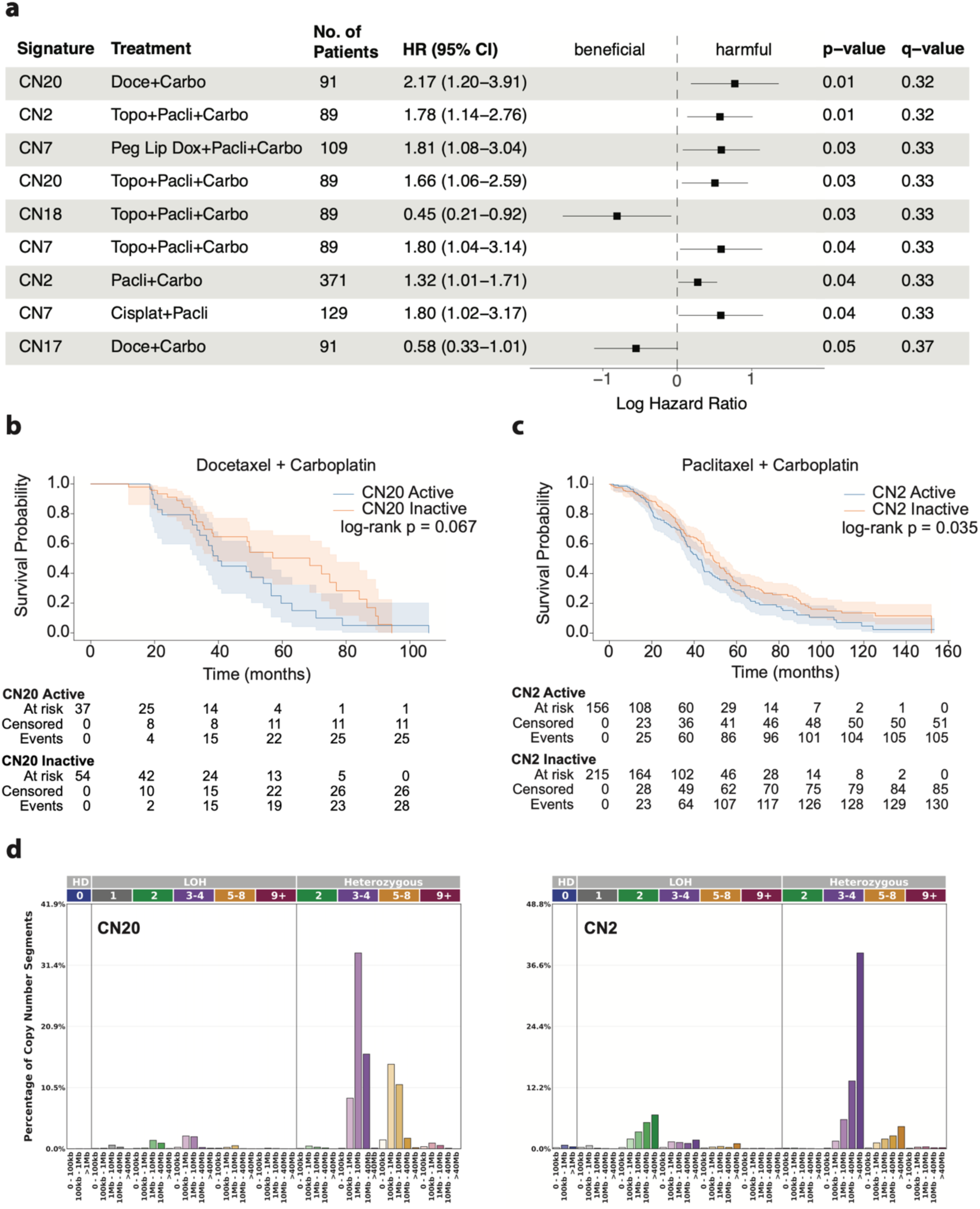
External evaluation of CN signature associations with overall survival in the GDC ovarian cancer cohort. **a**, Multivariate proportional hazard models showing the relative risk of CN signature exposures on OS for different therapies, controlling for age and tumor stage, in the GDC ovarian cancer cohort. The complete cohort comprised 485 patients. HRs represent the risk of any CN signature activity (CN signature activity > 0) compared to no activity (CN signature activity = 0). Treatments that were co-administered in >80% of cases were combined. The nine signature–treatment associations with the smallest p-values are displayed; q-values based on the Benjamini–Hochberg method. **b-c,** Kaplan-Meier curve comparing OS of **b**, CN20-active (CN20 signature activity > 0) and CN20-inactive (CN20 signature activity = 0) patients with ovarian cancer treated with Docetaxel + Carboplatin (n = 91) from the GDC cohort; **c**, CN2-active vs CN2-inactive patients with ovarian cancer treated with Paclitaxel + Carboplatin (n = 371). **d,** CN-category profiles of CN20 and CN2, showing the contributions of the 48 categories defined by loss-of-heterozygosity status, total copy-number state, and segment length.

## References

1. Alexandrov LB, Kim J, Haradhvala NJ, Huang MN, Tian Ng AW, Wu Y, et al. The repertoire of mutational signatures in human cancer. Nature. 2020;578(7793):94–101.

2. Steele CD, Abbasi A, Islam SA, Bowes AL, Khandekar A, Haase K, et al. Signatures of copy number alterations in human cancer. Nature. 2022;606(7916):984–91.

3. Macintyre G, Goranova TE, De Silva D, Ennis D, Piskorz AM, Eldridge M, et al. Copy number signatures and mutational processes in ovarian carcinoma. Nature genetics. 2018;50(9):1262–70.

4. Willing E-M, Vollbrecht C, Voessing C, Weist P, Schallenberg S, Herbst JM, et al. Development of the NOGGO GIS v1 assay, a comprehensive hybrid-capture-based NGS assay for therapeutic stratification of homologous repair deficiency driven tumors and clinical validation. Cancers. 2023;15(13):3445.

5. Borkar S, Markus F, Oetting A, Schmidt S, Vössing C, Horst D, et al. Detection of ESR1 mutations in tissue and liquid biopsy with novel next-generation sequencing and digital droplet PCR assays: insights from multi-center real life data of almost 6000 patients. Cancers. 2025;17(8):1266.

6. Moris C, Okrožnik N. cnakepit: A snakemake pipeline for copy number variant calling without normal tissue samples 2024 [Available from: https://github.com/pedricolino/cnakepit.]

7. Wang S, Li H, Song M, Tao Z, Wu T, He Z, et al. Copy number signature analysis tool and its application in prostate cancer reveals distinct mutational processes and clinical outcomes. PLoS Genetics. 2021;17(5):e1009557.

8. Monti S, Tamayo P, Mesirov J, Golub T. Consensus clustering: a resampling-based method for class discovery and visualization of gene expression microarray data. Machine learning. 2003;52(1):91–118.

9. Heath AP, Ferretti V, Agrawal S, An M, Angelakos JC, Arya R, et al. The NCI genomic data commons. Nature genetics. 2021;53(3):257–62.

10. Weinstein JN, Collisson EA, Mills GB, Shaw KR, Ozenberger BA, Ellrott K, et al. The cancer genome atlas pan-cancer analysis project. Nature genetics. 2013;45(10):1113–20.

11. Cao L, Huang C, Zhou DC, Hu Y, Lih TM, Savage SR, et al. Proteogenomic characterization of pancreatic ductal adenocarcinoma. Cell. 2021;184(19):5031–52. e26.

12. Islam SA, Díaz-Gay M, Wu Y, Barnes M, Vangara R, Bergstrom EN, et al. Uncovering novel mutational signatures by de novo extraction with SigProfilerExtractor. Cell genomics. 2022;2(11).

13. Zhao EY, Shen Y, Pleasance E, Kasaian K, Leelakumari S, Jones M, et al. Homologous recombination deficiency and platinum-based therapy outcomes in advanced breast cancer. Clinical Cancer Research. 2017;23(24):7521–30.

14. Joshi PM, Sutor SL, Huntoon CJ, Karnitz LM. Ovarian cancer-associated mutations disable catalytic activity of CDK12, a kinase that promotes homologous recombination repair and resistance to cisplatin and poly (ADP-ribose) polymerase inhibitors. Journal of Biological Chemistry. 2014;289(13):9247–53.

15. Etemadmoghadam D, Defazio A, Beroukhim R, Mermel C, George J, Getz G, et al. Integrated genome-wide DNA copy number and expression analysis identifies distinct mechanisms of primary chemoresistance in ovarian carcinomas. Clinical cancer research. 2009;15(4):1417–27.

16. Patch A-M, Christie EL, Etemadmoghadam D, Garsed DW, George J, Fereday S, et al. Whole–genome characterization of chemoresistant ovarian cancer. Nature. 2015;521(7553):489–94.

17. Sztupinszki Z, Diossy M, Krzystanek M, Reiniger L, Csabai I, Favero F, et al. Migrating the SNP array-based homologous recombination deficiency measures to next generation sequencing data of breast cancer. NPJ breast cancer. 2018;4(1):16.

18. German Cancer Society, German Cancer Aid, AWMF. S3 guideline: Diagnosis, therapy and follow-up of malignant ovarian tumors 2024 [Available from: https://register.awmf.org/de/leitlinien/detail/032-035OL.]

19. Maclachlan KH, Rustad EH, Derkach A, Zheng-Lin B, Yellapantula V, Diamond B, et al. Copy number signatures predict chromothripsis and clinical outcomes in newly diagnosed multiple myeloma. Nature communications. 2021;12(1):5172.

20. Drews RM, Hernando B, Tarabichi M, Haase K, Lesluyes T, Smith PS, et al. A pan-cancer compendium of chromosomal instability. Nature. 2022;606(7916):976–83.

21. Spruck CH, Won K-A, Reed SI. Deregulated cyclin E induces chromosome instability. Nature. 1999;401(6750):297–300.

22. Network CGAR. Integrated genomic analyses of ovarian carcinoma. Nature. 2011;474(7353):609.

23. Ciriello G, Cerami E, Sander C, Schultz N. Mutual exclusivity analysis identifies oncogenic network modules. Genome research. 2012;22(2):398–406.

24. Fukasawa K. Centrosome amplification, chromosome instability and cancer development. Cancer letters. 2005;230(1):6–19.

25. Bagheri-Yarmand R, Biernacka A, Hunt K, Keyomarsi K. Low Molecular–Weight Cyclin E Overexpression Shortens Mitosis, Leading to Chromosome Missegregation and Centrosome Amplification. Cancer research. 2010;70(12):5074.

26. Xu H, George E, Gallo D, Medvedev S, Wang X, Datta A, et al. Targeting CCNE1 amplified ovarian and endometrial cancers by combined inhibition of PKMYT1 and ATR. Nature Communications. 2025;16(1):3112.

27. Farooq AR, Zhang AX, Chan-Seng-Yue M, Topham JT, O’Kane GM, Jang GH, et al. The tandem duplicator phenotype may be a novel targetable subgroup in pancreatic cancer. NPJ Precision Oncology. 2025;9(1):100.

28. Menghi F, Barthel FP, Yadav V, Tang M, Ji B, Tang Z, et al. The tandem duplicator phenotype is a prevalent genome-wide cancer configuration driven by distinct gene mutations. Cancer cell. 2018;34(2):197–210. e5.

29. Davies H, Glodzik D, Morganella S, Yates LR, Staaf J, Zou X, et al. HRDetect is a predictor of BRCA1 and BRCA2 deficiency based on mutational signatures. Nature medicine. 2017;23(4):517–25.

30. Dubois FP, Shapira O, Greenwald NF, Zack T, Wala J, Tsai JW, et al. Structural variants shape driver combinations and outcomes in pediatric high-grade glioma. Nature cancer. 2022;3(8):994–1011.

31. Thompson JS, Madrid L, Hernando B, Sauer CM, Vias M, Escobar-Rey M, et al. Predicting resistance to chemotherapy using chromosomal instability signatures. Nature Genetics. 2025;57(7):1708–17.

